# Probing the Structural Basis of Citrus Phytochrome B using Computational Modelling and Molecular Dynamics Simulation Approaches

**DOI:** 10.1101/2021.05.11.443630

**Authors:** Muhammad Tahir ul Qamar, Muhammad Usman Mirza, Jia-Ming Song, Muhammad Junaid Rao, Xitong Zhu, Ling-Ling Chen

## Abstract

Phytochromes (Phys) are known as red/far-red light photoreceptors and are responsible for directing the photosensory responses across the species, majorly from fungal, bacterial and plant kingdoms. Such responses majorly include photosynthetic potential and pigmentation in bacteria, whereas in a plant, they are involved in chloroplast development and photomorphogenesis. Many prokaryotic Phys have been modelled for their structural and functional analysis, but their plant counterparts have not been explored yet. To date, only the crystal structures of the photo-sensing module (PSM) of PhyB isoform from *Arabidopsis thaliana* and *Glycine max* have been resolved experimentally. Thus, in this study, we elucidated the complete 3D structure of Citrus PhyB. Initially, the structure and organisation of the Citrus PhyB have been predicted computationally, which were found to have the same domain organisation as *A. thaliana* and *G. max* PhyBs, yet their considerable distinct structural difference indicated potential divergence in signaling and functioning. Therefore, to evaluate the structural and functional implications of Citrus PhyB, we compared its structure with *A. thaliana* and *G. max* PhyBs using molecular dynamics (MD) simulation approaches. The modeling studies revealed that the region of Citrus PhyB-GAF domain possibly contributes to the variations between Citrus, *A. thaliana* and *G. max* PhyBs structures/functions. Hence, structural and molecular insights into Citrus PhyB can help to discover the Phys signaling and thus, an essential framework can be designed for optogenetic reagents and various agricultural/horticulture benefits.

**One sentence summary:** A complete Citrus PhyB structure together with photo-sensory and out-put modules provides significant information to evaluate its biological activities for agricultural benefits.

## INTRODUCTION

Among photoreceptors, plant phytochromes are known to make a huge contribution towards environmental development changes occurring in nature that may include germination, flowering, and de-etiolation. Around 20% of genes responsible for these events are known to be regulated by particular phytochromes regulating light-dependent processes (Song et al., 2018). Phytochromes have a unique attribute to reversibly interconvert between dark-adapted and photoactivated end states, thus regulating a wide array of light-dependent processes (Burgie and Vierstra, 2014; Burgie et al., 2016). Two stable photo-interconvertible states adapted by phytochromes include a red light-absorbing Pr state and a far-red light-absorbing Pfr state (Quail, 2002; Ulijasz and Vierstra, 2011). Initially, they are produced as Pr, the inactive ground state, which converts to Pfr (biologically active state) only upon red light irradiation (Ulijasz and Vierstra, 2011). As a reversible process, far-red irradiation of Pfr can then convert back the phytochrome to Pr, which is the only reversible ‘photo-switches’ mechanism known in nature. Heme-derived bilin (or linear tetrapyrrole) phytochromobilin (PΦB) is the key responsible agent for this unique photochemistry and gets covalently attached to the protein moiety, forming cysteine thioether linkage (Rockwell et al., 2006; Ulijasz and Vierstra, 2011). Several researches over past years now have revealed that higher plants usually possess many isoforms of Phys which play crucial roles in almost all aspects of a plant’s life, majorly including seed germination, fruit ripening, flowering time, shade avoidance, photoperiodism, and photomorphogenesis (Ulijasz and Vierstra, 2011).

Overall phytochrome family has been revealed with a homodimer modular architecture after careful biochemical and sequence analyses, where each subunit is divided into PSM and OPM modules (Li et al., 2010; Burgie et al., 2014). On Phys N-terminal, there lies a PSM module containing Per/Arndt/Sim (PAS), cGMP phosphodiesterase/adenylyl cyclase/FhlA (GAF), and Phytochrome (PHY) domains (Rockwell et al., 2006). The GAF domain having intrinsic bilin-ligase activity also provides a binding pocket for non-covalent binding of PΦB along with cysteine linkage. Along with it, PAS and PHY domains also play a critical role to maintain stabilised Pfr conformation. Usually, the length of the N-terminal varies among plant phytochromes that may be responsible for their altered biological activities and stable conformations. Likewise, it has been reported that the C-terminal region of Phys possesses an OPM module that has adjacent PAS, PAS, and histidine kinase-related (HKR) domains which can control various signaling cascades upon Pfr formation (Li et al., 2010; Burgie et al., 2014).

Among various Phys present in plants, three discrete apoprotein-encoding genes (PhyA– PhyC) are conserved within angiosperms (Mathews et al., 1995; Franklin and Quail, 2010). Some different Phys have also been identified in dicotyledonous plants, which are considered to be related to gene duplication events (Mathews et al., 1995; Li et al., 2015). Only five genes (PhyA–PhyE) encoding phytochrome apo-proteins have been retrieved, characterised and sequenced in the model species *A. thaliana* (Clack et al., 1994; Franklin and Quail, 2010). Among them, PhyA being a primary photoreceptor, is known as the controlling agent of high-irradiance FR light responses, including cotyledon expansion, hypocotyl elongation and seed germination (Reed et al., 1993). Moreover, PhyB is another photoreceptor primarily of red-light and thus known to mediate shade-avoidance syndrome responses. Such responses result in internode elongation, reduced leaf area, petiole elongation and early flowering (Reed et al., 1993; Sheehan et al., 2004). The PhyB protein is also a homolog of PhyD (having 80% sequence similarity) and forms a distinct evolutionary subgroup with PhyE (Goosey et al., 1997; Franklin and Quail, 2010). Though, both of them are not functionally redundant (Sharrock et al., 2003; Sheehan et al., 2004) and majorly control the flowering time along with petiole and internode elongation. Similarly, PhyC is responsible for generating seedling response against red light and also suppresses flowering under short-day conditions (Franklin et al., 2003; Monte et al., 2003; Sheehan et al., 2004).

The present paper focuses on structural insights of Citrus PhyB. Similar to *A. thaliana* (Sharrock and Clack, 2002) and tomato (Hauser et al., 1995), Citrus genome also encodes five phytochromes (PhyA-PhyE). Little is known about PhyB functions in plants. Most of the studies have focused on its roles in the integration of light and temperature signals (Legris et al., 2016), reduced irradiance of canopy shades (Trupkin et al., 2014) and growth/defense (Campos et al., 2016). Though, the structural conservation of PhyB is yet unclear as significant difference has been observed among sequence and domain architecture. Therefore, its particular characteristics and action mechanism can be studied, which is of great interest. Hence, in the current study, the full-length structure of Citrus PhyB in apo/holo states has been explored employing protein-structure modelling along with molecular docking and MD simulation (Tahir Ul Qamar and Khan, 2017; Maryam et al., 2019; Khalid et al., 2020; Maryam et al., 2020). Initially, the three-dimensional (3D) structure of Citrus PhyB was predicted and docking of phytochromobilin ligand was evaluated with its GAF domain. To analyse the stability and conformational transitions of the complex over time, 100 ns MD simulations were performed. Overall, the study has been aimed to get insights into plant Phys signaling and discovering essential structural information of Citrus PhyB to evaluate its biological activities for agricultural/horticulture benefits.

## RESULTS AND DISCUSSION

PhyB eminent by a long glycine/serine-rich N-terminal extension is one of the most important and active members among phytochrome family. Previous efforts to obtain full-length atomic structure of PhyB in plants have not been successful. Only PSM-PhyB modules of *A. thaliana* (Pettersen et al., 2004) and *G. max* (Nagano et al., 2020) have been reported. In this study, we probe and explore the domain-to-domain structural orientation of complete Citrus PhyB using computational and biophysics approaches, to understand its molecular mechanism. Citrus PhyB sequence is composed of 1137 residues and likely a homodimer similar to *A. thaliana* (Pettersen et al., 2004) containing two sections of polypeptide chains, including an N-terminal photosensory module (PSM) and an output module (OPM) towards the C-terminal. The PSM harbors consecutively Per/Arnt/Sim (PAS) domain (85—201), cGMP phosphodiesterase/adenylyl cyclase/FhlA (GAF) domain (234—422), and a Phy-specific (PHY) domain (425—598). The OPM sequentially contains two adjacent PAS domains (PAS1: 639— 745 and PAS2: 761—880) and histidine kinase-related (HKR) domain (909—1129) (Figure 1A). Secondary structure of PhyB is based on 20% alpha helixes, 10% extended strands and 30% random coils (supplemental figure 1). A protein with more helices is more flexible to fold and likely to interact with other proteins (Tokuriki and Tawfik, 2009).

**Figure 1.**
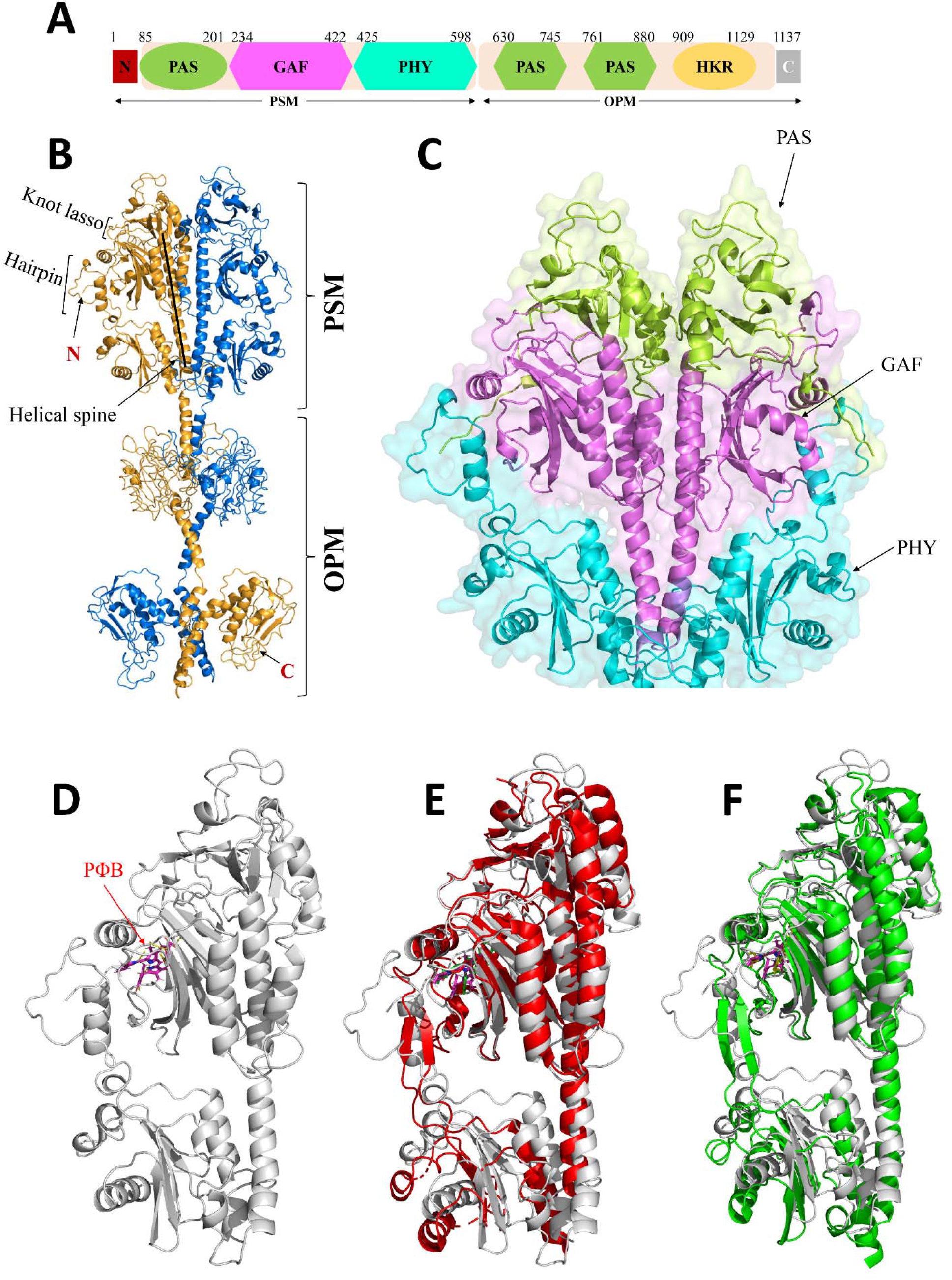
Structural basis of Citrus PhyB. (A) Basic structure and domain boundaries of PSM and OPM modules of Citrus PhyB. (B) Front side view of 3D cartoon structures of complete Citrus PhyB homodimer in apo form. Both chains are highlighted with different colors, and knot lasso, hairpin region and helical spine are highlighted. (C) Close up view of apo-PSM module in homodimer form. The PAS, GAF and PHY domains are colored in green, pink and cyan, respectively. (D) Close up view of holo-PSM module in monomer form. The attached phytochromobilin (PΦB) is highlighted with pink color. (E) Superimposition of Citrus (grey) and *A. thaliana* (red) holo-PSM modules in monomeric states. (F) Superimposition of Citrus (grey) and *G. max* (green) holo-PSM modules in monomeric states.

### Structural elucidation of PhyB

With the advent of computational technology, there is a massive increase in sequence submission in various databases, but only 1% of these submitted sequences have been evaluated and annotated experimentally (Tahir Ul Qamar and Khan, 2017). Proper structural analysis of protein can aid functional characterisation of proteins and has been used in various studies to determine the structure and function of several proteins. Sequence-to-structure alignment of Citrus PhyB with the crystal structures of PhyB-PSMs of *A. thaliana* and *G. max* revealed 46% query-coverage each, 0.00 E-value each, and 85.42% and 86.23% sequence identity, respectively (supplemental figure 2). Since the query-coverage was low, domain-to-domain modeling of complete PhyB was performed to predict its full-length structure covering both PSM and OPM modules.

For accurate modeling, all individual domains homology/comparative models of Citrus PhyB were built first. RMSD of all generated domain models were found between 0.1-0.3 Å. Ramachandran’s analysis revealed that 98-99% residues of all predicted domain models were in favorable regions. Individual domains were then combined into their corresponding PSM and OPM modules apo/holo monomeric structures, and later individual monomeric chains were modeled into a complete homo-dimeric 3D structure as shown in Figure 1. Both polypeptide chains were examined for their secondary structure integrity and interactions with each other to estimate the stability of Citrus PhyB homodimer sate. Results revealed that the secondary structure of both polypeptide chains is intact, and established 76 hydrogen bonds and 7 salt bridges in total with each-other (supplemental table 1), which displays the high stability of PhyB homodimer and support the hypothesis that Citrus PhyB is a homodimer similar to *A. thaliana* (Pettersen et al., 2004).

The overall molecular architecture of Citrus PhyB homodimer showed a parallel head-to-head arrangement of sister chains interceded through two juxtaposed α-helices. The PAS, GAF and PHY domains were placed in the PSM module and PAS1, PAS2 and HKR domains were placed in OPM module. Not surprisingly, the predicted Citrus PhyB model revealed the domain organisation is similar with the PSM crystal structures of *A. thaliana* (PDB ID: 4OUR) and *G. max* (PDB ID: 6TL4) (Nagano et al., 2020) with 4.195 Å and 5.004 Å RMSD differences, respectively, and slight shifts in domain positions (Figure 1E-F). Significant differences were observed towards the C-terminal of all three PSM structures, especially in PHY domain regions. However, the GAF domain binding pocket was found remarkably similar to *A. thaliana* and *G. max* pockets. The C-terminal region of PSM is comprised of PHY overlaps with the N-terminal region (PAS1) of OPM module. Full-length Citrus PhyB domain assembly revealed that the catalytic end containing HKR domain is rotated around the dyad axis, which has orientated HKR domain opposite to the rest of other domains in the individual chains. The structure was further optimised for energy minimisation and to add missing residues, heavy atoms, polar hydrogen and charges. Quality analysis of final full-length Citrus PhyB revealed 98.4% residues are in the favorable regions, whereas only 1.6% residues were found in the outlier region, as shown in supplemental figure 3.

It has been well established that phytochromes switch between Pr and Pfr states and PΦB is the key responsible agent for this unique mechanism. The GAF domain has been extensively characterised in several studies (Essen et al., 2008; Li et al., 2010; Burgie et al., 2014), therefore it can provide a solid reference background to get deep insights into the conformational stability of Citrus PhyB. The GAF domain binding pocket of Citrus PhyB (supplemental figure 4) was explored further, and it was noticed it shares strong homology with previously published phytochromes structures, including *Sorghum bicolor* PhyB-GAF domain (PDB ID: 6TC5) (Nagano et al., 2020). Although structural differences were observed, but bounded PΦB adapted confirmation in Citrus GAF binding pocket similar to *A. thaliana, G. max* and *S. bicolor* co-crystallised PΦB binding confirmations, and thus validating the holo structure of Citrus PhyB (Figure 2). Only 1.669 Å RMSD difference was observed between Citrus and *S. bicolor* GAF domains structures. The PΦB was attached to Citrus GAF domain by making several hydrogen bonds and van der Waals interactions. It made hydrogen bonds with important binding site residues Asp289, Arg334, His340, Ser352 and His382. While, it made polar interactions with Met256, Tyr258, Tyr280, Leu283, Ile290, Pro291, Ser294, Phe298, Arg334, Cys339, His337, Tyr343 and Ala354. Previous studies reported that these residues play a critical role in keeping the binding pocket integrity and formation of PΦB and GAF domain complex (Essen et al., 2008; Li et al., 2010; Burgie et al., 2014; Nagano et al., 2020).

**Figure 2.**
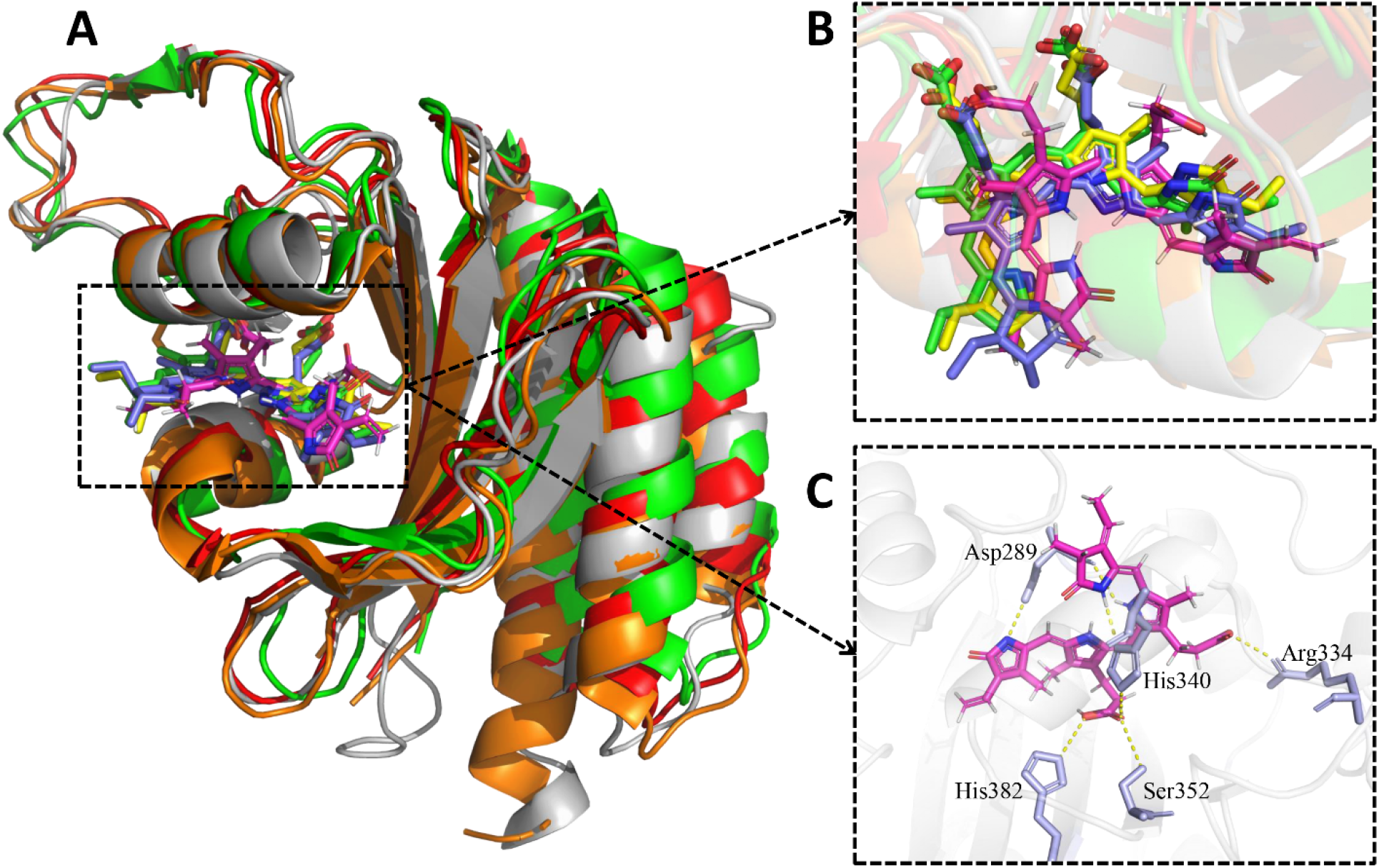
The GAF domain of Citrus PhyB. (A) Superimposition state close-up view of Citrus (grey), *A. thaliana* (red), *G. max* (green) and *S. bicolor* (orange), PΦB bound GAF domains. (B) Superimposition of co-crystalized PΦB with *A. thaliana* (green), *G. max* (yellow) and *S. bicolor* (blue) and docked pose of PΦB with Citrus (pink), GAF domains of PhyB. (C) Molecular interactions of PΦB with hot-spot residues of Citrus PhyB-GAF domain.

### MD simulation analysis

Characterizing the backbone dynamics of multi-domain proteins in terms of residual fluctuations and associated motions using long-run MD simulations is a practical and fundamental step towards interpreting molecular behaviour and mechanism of regulation of multidomain proteins. MD simulation has gradually become a crucial approach to understanding proteins structures’ physical basis and their important biological functions (Patodia et al., 2014). To evaluate the conformational stability and analyze different functions of PhyB in Citrus, PSM module was selected to study through MD simulation analyses. PSM module of PhyB has been extensively characterized in several studies (Essen et al., 2008; Li et al., 2010; Burgie et al., 2014; Nagano et al., 2020), therefore it can provide a solid reference background to get deep insights into conformational stability of Citrus PhyB. Crystal structures of *A. thaliana* (PDB ID: 4OUR) and *G. max* (PDB ID: 6TL4) PhyB-PSM modules bound to PΦB were used as references. To analyze the structural stability and dynamics, several parameters including, RMSD, Rg and RMSF scores were evaluated by CPPTRAJ in Amber16 using 100ns MD simulations (Roe and Cheatham III, 2013).

### RMSD analysis

RMSD is a standard measure of structural distance between a group of atoms and principally used to compute the backbone variation compared with its original structural conformation. In this context, the dynamic stability of a protein can be determined by the variations observed in protein’s backbone for the period of simulations. The stability of protein structure can be determined over a simulation period, which is reflected by the convergence of RMSD trajectory over time and hence, represents the most favorable conformation. Here in, RMSD of the holo *A. thaliana, G. max* and Citrus, and apo Citrus PhyB-PSM systems were analyzed for all C_α_ backbone atoms, and the results reveal significant variations as compared to their initial conformations. RMSD trajectories of all 4 systems for 100 ns are shown in Figure 3. Overall, the average RMSD values for *A. thaliana, G. max*, holo-Citrus and apo-Citrus were 4.04 Å, 4.11 Å, 4.84 Å and 5.19 Å, respectively. Both holo- and apo-Citrus PSM systems displayed overall minor fluctuations than *A. thaliana* and *G. max*. Among these, the holo-Citrus PSM found to be relatively more stable as compared to its apo conformation, which was evident due to its ligand-bound conformation, that allowed consistent hydrogen bond interactions over time and represented a favourable conformation with PΦB. Whereas, the average RMSD value of apo-Citrus system was higher than the other three systems due to the unbound conformation of protein and underwent more flexible over time. The higher RMSD at some intervals during simulation of holo systems is probably associated with the unbinding and binding of some unstable interactions in the GAF binding pockets of PSM modules. These results showed that overall Citrus PSM is stable. At the same time, some of its regions possess variations in its apo form, which regions may be responsible for the distinct functions of Citrus PhyB.

### Rg analysis

Protein conformational stability can also be analyzed using the Rg which monitors the effective size of a particular protein and then measures its compactness (Khalid et al., 2018). Rg is actually the mass-weighted root mean square distance of a group of atoms from their common center of mass. The Rg analysis of proteins also coincides with the RMSD stability analysis performed in this study (Figure 3). Similar to RMSD, all systems displayed minor to major fluctuations throughout the 100 ns simulation period. However, all systems remained compact with average Rg value for the *A. thaliana* 27.94 Å, for *G. max* 28.90 Å, for holo-Citrus 27.90 Å and for apo-Citrus 28.95 Å. The Rg value for apo-Citrus PSM system remained higher compared to other systems under study. While the average value for holo-Citrus PSM system remained lower compared to *A. thaliana* and *G. max*. The fluctuations in Rg value at different intervals in systems is probably because of unbinding and binding of PΦB with PSM modules. These results demonstrated that the PΦB binding contributed towards the compactness of the systems over the 100 ns simulation time.

**Figure 3.**
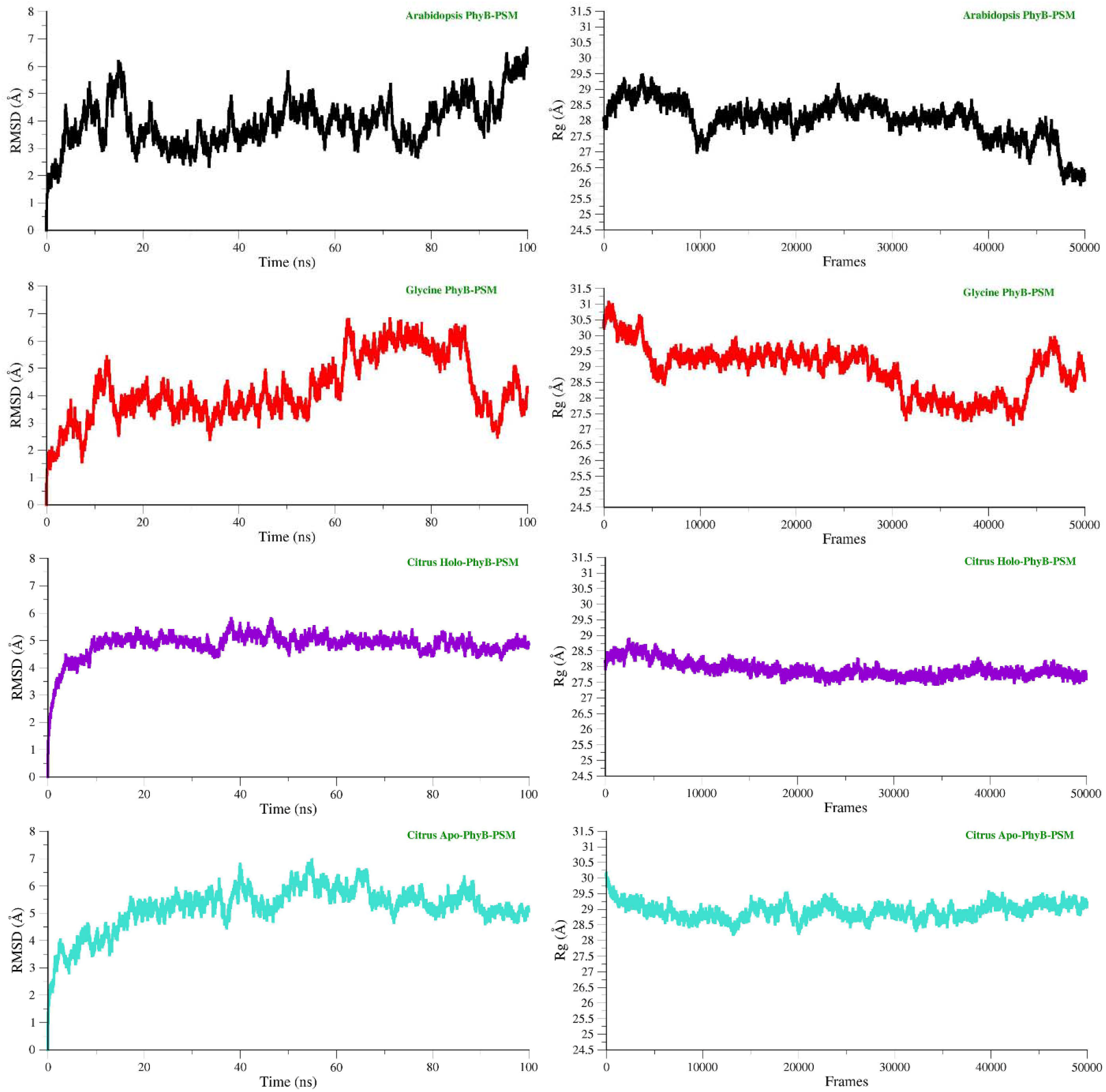
The schematic illustration of RMSD and Rg results of all the simulated systems. The RMSD and Rg of *A. thaliana, G. max*, holo Citrus and apo Citrus systems are represented in black, red, purple and cyan colors, respectively.

### RMSF analysis

To investigate the mobility of residues influenced by PΦB and to provide deep insights into the dynamics behavior of all systems due to the binding/unbinding of ligand, RMSF stability analysis was performed (Khalid et al., 2018), which measures the average deviation of a protein residue over time from its reference position. The obtained peaks represent the areas of increased residual flexibility and describe the influence upon ligand’s binding. Flexibility and rigidity play essential roles in different biological processes. In the case of *A. thaliana* and holo-Citrus, higher fluctuations were observed, especially in the start and between 314 to 410 amino acids within GAF domain region. While *G. max* and apo-Citrus systems showed relatively lesser fluctuations than other two systems (Figure 4A). The average RMSF values for *A. thaliana, G. max*, holo-Citrus and apo-Citrus were 4.03 Å, 1.98 Å, 4.59 Å and 1.95 Å, respectively. Subsequently, we further analyzed the RMSF of GAF domain of each system (Figure 4B). Major fluctuations were observed in three sheets (β11, β12, β13), five loops (γ20, γ21, γ22, γ23, γ24) and 3 helixes (α9, α10, α11) of Citrus systems, which are core parts of GAF binding pocket and vital for PΦB binding. These larger fluctuations revealed increased random motions of residues located in these particular regions. Fluctuations in the GAF binding region were considerably higher than the others, which implies that this might be due to the important residues that are vital for PΦB binding. These results suggested that the GAF region (314-410 residues) can possibly contribute to the variations between Citrus, *A. thaliana* and *G. max* PhyB structures and functions.

**Figure 4.**
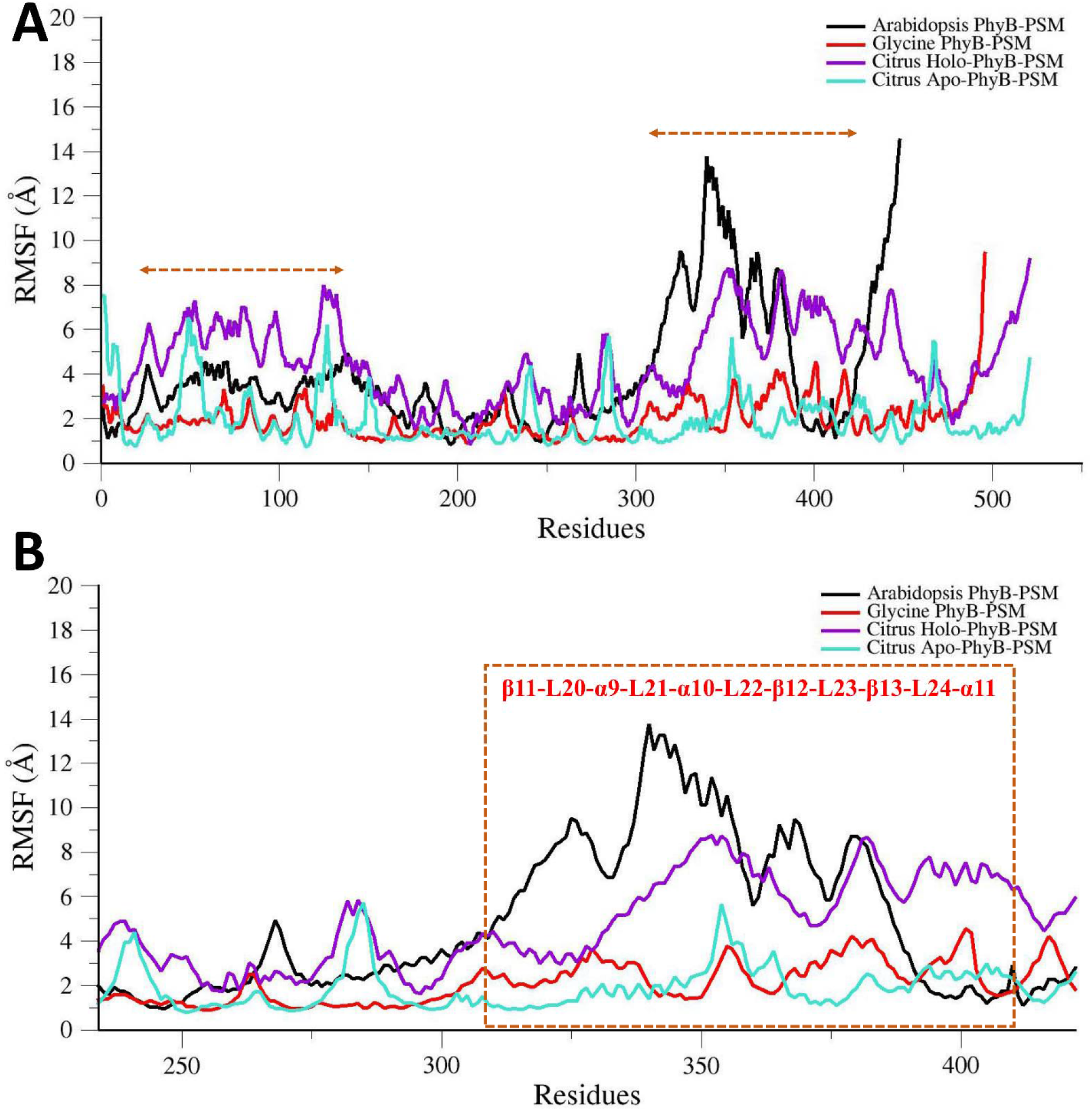
The schematic illustration of residual flexibility index of *A. thaliana, G. max*, holo Citrus and apo Citrus systems over the period of 100 ns simulation. (A) RMSF plot of all systems, where highly fluctuated regions are highlighted with arrows. (B) RMSF plot of GAF regions residues. The highly fluctuated and important secondary structures for PΦB binding are highlighted in the box.

### Principal component analysis (PCA)

To further investigate the PΦB binding dynamics with GAF domain and connect protein conformational changes within the GAF binding domain with the binding of PΦB, PCA analysis was performed. Since, during the RMSF analysis holo-Citrus system showed more similar behavior like holo *A. thaliana* system, thus *A. thaliana* system was used as a reference to identify the overall normalized pattern of motion. Functionally critical structural transitions can be revealed by analyzing the diagonalization of the covariance matrix, which can then infer the meaningful configurational space (Khalid et al., 2018). The essential dynamics were performed for the backbone atoms of *A. thaliana* and holo-Citrus GAF domains. Total 87% of the overall positional transitions were found to be corresponded by the initial principal components in all the systems, and their motion was also examined via normal mode analysis in VMD, and the porcupine plot was generated to evaluate the magnitude and direction of selected eigenvectors (Tai et al., 2002). Regions of Citrus GAF domain contributing to high variance during essential dynamic analysis are shown in Figure 5. The regions include a loop (γ23) starting from GAF domain sequence 360 to 373 amino acids, which has also been indicated by the residual stability analysis. Thus, predicting that this region can contribute to the variations between Citrus and *A. thaliana* PhyB functions.

**Figure 5.**
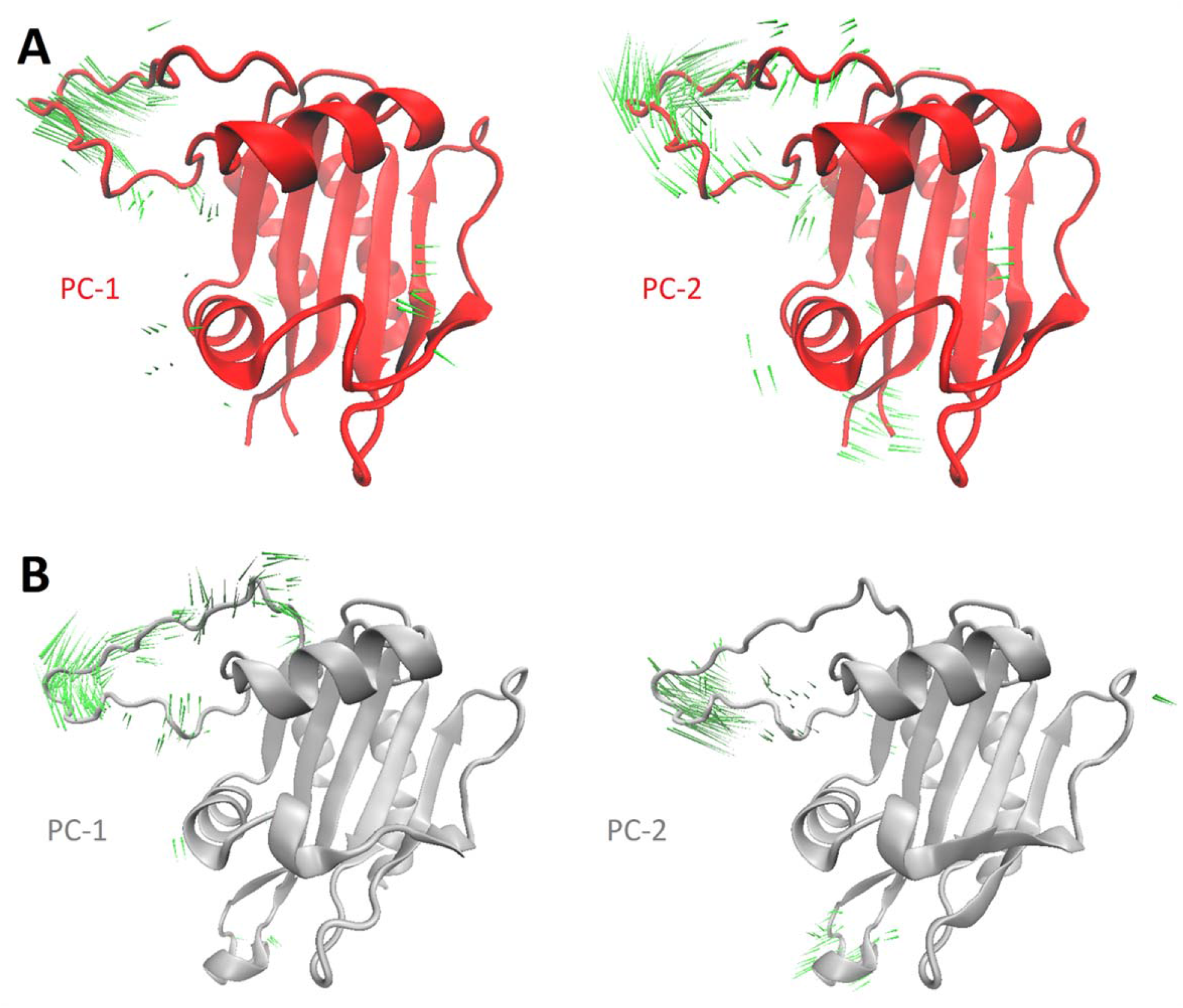
Porcupine plots illustrating the highest principal component variance of the first (PC-1) and second (PC-2) modes of motion, estimated through uncorrelated trajectories analysis. (A) Holo A. thaliana GAF domain, (B) Holo-Citrus GAF domain. Green color porcupines showing the regions (γ) with significant movement, and their length represents the degree of mobility.

### Binding free energy calculations using MM/GBSA method

To further assess the affinity and interaction energies changes of the PΦB bound PSM modules of *A. thaliana, G. max* and Citrus, we performed MM/GBSA analysis (Genheden and Ryde, 2015). The MM/GBSA analysis results confirmed a favorable affinity of PΦB within the binding pocket located in the GAF domain of the PSM module **(Table 1)**. The binding free energies calculated by MM/GBSA are more accurate end-point method for analyzing the docked complexes. In this method, the results also highlight the electrostatic, Van der Waals (vdW), and non-polar solvation energy terms favorably contribute towards the binding energy. Since van der Waals represents the strength of nonpolar interactions derived as a result of binding between ligand and binding pocket, it was estimated to have the highest contribution towards the binding energy with comparatively more negative values. Besides, polar solvation energy represents the energy associated with dissolving PΦB within the solvent. Highly positive polar solvation energies were obtained against all three PhyB-PSMs, depicting their preference for a hydrophobic environment.

**Table 1.**
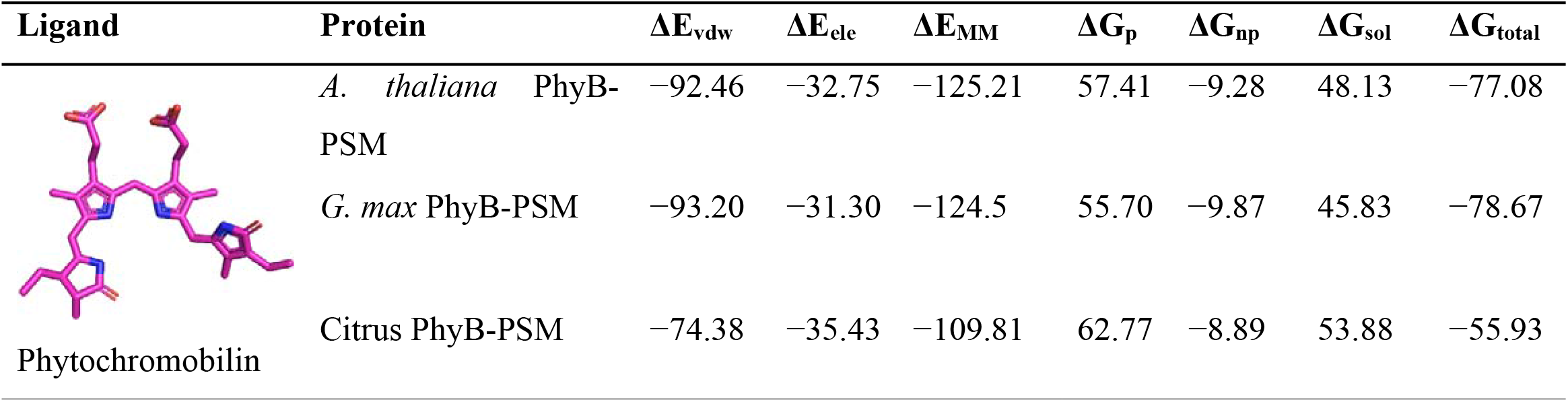
Binding free energy calculations of PΦB bound *A. thaliana, G. max* and Citrus PhyB-PSMs. All energy values are calculated in kcal/mol.

| Ligand | Protein | $\Delta E_{vdw}$ | $\Delta E_{ele}$ | $\Delta E_{MM}$ | $\Delta G_p$ | $\Delta G_{np}$ | $\Delta G_{sol}$ | $\Delta G_{total}$ |
| --- | --- | --- | --- | --- | --- | --- | --- | --- |
| Phytochromobilin | *A. thaliana* PhyB-PSM | -92.46 | -32.75 | -125.21 | 57.41 | -9.28 | 48.13 | -77.08 |
|  | *G. max* PhyB-PSM | -93.20 | -31.30 | -124.5 | 55.70 | -9.87 | 45.83 | -78.67 |
|  | Citrus PhyB-PSM | -74.38 | -35.43 | -109.81 | 62.77 | -8.89 | 53.88 | -55.93 |

The comparative results represent highly favorable Van der Waals energies for PΦB. Considering that electrostatic interactions with surrounding residues play a marginal role, indicating the complex’s stability depends on the hydrophobic contacts established within the pocket. It has been found that these electrostatic interactions are important for the stability of PΦB, although the hydrophobic interactions still play a significant role in their binding affinity. From the calculated results, it can be noted that ΔG_total_ of PΦB was low against Citrus PhyB-PSM (−55.93 kcal/mol), as compared to *A. thaliana* PhyB-PSM (−77.08 kcal/mol) and *G. max* PhyB-PSM (−78.67 kcal/mol). Whereas all systems displayed significant high negative values for vdW interaction energies against Citrus PhyB-PSM (−74.38 kcal/mol), *A. thaliana*-PSM (−92.46 kcal/mol) and *G. max* PhyB-PSM (−93.20 kcal/mol).

After examining the binding mode and hydrogen bond occupancy of PΦB against Citrus PhyB-GAF (Figure 2C), it was noteworthy to highlight the independent contribution of residues within GAF binding pocket towards the binding energy per-residue energy decomposition analysis (Genheden and Ryde, 2015) was carried out for PΦB bound Citrus PhyB-PSM system. Binding free energy decomposition analysis reveals the contribution of each amino acid’s sidechain and backbone atoms. Studies have illustrated the use of binding free energy calculations and its per-residue decomposition methods towards understanding the mechanisms of protein-protein binding as well as binding of protein-ligand complexes (Gohlke et al., 2003; Xu et al., 2013; Genheden and Ryde, 2015; Sun et al., 2018). Per-residue decomposition analysis also contributed towards dynamically estimating the interactions of compounds within the binding pocket. The results indicated that PΦB could establish from 4 to 6 medium-to high-affinity interactions with residues belonging to Citrus GAF domain alongside interacting with surrounding residues of the binding site. The analysis of residual energy contribution (ΔG_residue_) to the total binding free energy of PΦB against Citrus PhyB revealed that residues Asp289 (−2.34 kcal/mol), Arg334 (−2.61 kcal/mol), His340 (−3.17 kcal/mol), Ser352 (−2.11 kcal/mol), His382 (−3.24 kcal/mol), Cys339 (−1.41 kcal/mol) and Thr88 (−1.19 kcal/mol) were found to contribute the most (ΔG_residue_ < −1 kcal/ mol) towards the binding of PΦB to Citrus PhyB. Although these potential residues were located in the region with major fluctuations (Figure 4), the consistent interactions with the PΦB binding sampled more favorable conformation over a simulation period.

These results from MM/GBSA binding free energy and per residue decomposition analysis strengthen the hypothesis that PΦB may anchor between residues of the cavity and surrounding regions in Citrus PhyB, which may be responsible for the distinct functions of Citrus PhyB. These interactions remained consistent throughout simulation as indicated in the RMSD plot of PΦB bound Citrus, which remained stable compared to *A. thaliana* and *G. max* (Figure 3). Overall, the MD simulations and MM/GBSA calculations indicated a representative binding mode highlighting highly stable interactions of PΦB against Citrus PhyB.

## CONCLUSION

Phytochromes, being known as red/far-red light photoreceptors of plants, also play several significant roles in transcriptional regulation of genes, initiation of germination/flowering, and controlling the developmental stages of plants. Although known for many years, their key fundamental features are still poorly understood. Till now, the crystal structure of only PSM module of *A. thaliana* and *G. max* have been resolved, and efforts to resolve complete PhyB have not yet been successful. Therefore, PhyB of Citrus was further studied for its structure and function by using *A. thaliana* and *G. max* PhyBs as reference. It was found that Citrus PhyB has significant structural dynamics variations as compared with *A. thaliana* and *G. max* PhyBs. Its GAF domain, knot region, and helical spine show distinct structural differences potentially important to signaling. Moreover, the structural elucidation of PSM modules of *A. thaliana, G. max* and Citrus PhyBs through MD directed energy calculations significantly contributed additional value in overall comparison. In conclusion, the details about Citrus PhyB structure given in the present study will enable molecular insights into plant Phys signaling, and provide essential structural information to redesign their activities for agricultural/horticulture benefits.

## MATERIALS AND METHODS

### Computational modeling of complete PhyB

There is no experimentally solved full-length structure of the plant’s PhyB, and efforts to obtain a complete atomic structure of PhyB have not yet been productive. To define the structure of Citrus PhyB, its amino acids sequence (ID: Cs9g02220.1) was retrieved from our *Citrus sinensis* annotation project database (http://citrus.hzau.edu.cn/orange/index.php) (Wang et al., 2014) and subjected to PSI-BLAST search against the Protein Data Bank (PDB). All the templates identified in the BLAST search were < 46% in query coverage to the query sequence and mostly covering the PSM part of PhyB sequence. In the absence of an appropriate template covering the full-length sequence of Citrus PhyB, and to understand accurate molecular architecture of full-length PhyB, individual domain models were initially predicted through Modeller v 9.20 (Webb and Sali, 2016). For all individual domain models of PhyB, PSI-BLAST was employed at first to identify homologous templates from Protein Data bank (PDB) (Berman et al., 2000). In the next step, sequence to structure alignment of the query sequence and template structures was done in Fugue (Miguel et al., 2002). Fugue is a freely available distant homolog identification tool, which identifies the extent of conservation and variation between the query sequence and template structures (Shi et al., 2001). Fugue is also integrated into the Modsuite pipeline (Unpublished Skwark MJ, Ochoa-Montaño B and Blundell TL) (Maryam et al., 2019). Further, in the homology modeling, Fugue generated alignment was employed in a freely available JOY tool to generate a sophisticated graphical representation and to identify the structural features of the query protein sequence, secondary structure composition, and ligand binding sites (Mizuguchi et al., 1998; Miguel et al., 2002). Based on the fugue alignment, the Modeller v9.20 tool was then directed to performs homology modeling of query sequences (Webb and Sali, 2017). Initial models of PAS, GAF, PHY, and HKR domains were selected based on the Discrete Optimized Protein Energy (DOPE) score (Shen and Sali, 2006) and GA341 values (Eswar et al., 2008). Rigorous energy minimisation of the initial models was performed to relieve steric clashes between the side/main-chain atoms of the adjacent residues.

Multi-domain proteins’ modeling where all the respective domains are placed in accurate relative orientation is among the challenging tasks of structural biology. The Phys are well known for their photo-switching properties, and they can switch between Pr (in-active/apo) and Pfr (active/holo) states. Structural insights, including conformational heterogeneity at the 3D structural level, are crucial to decipher the regulation of PhyB in these states. In this context, after building individual domain models, N-terminal PSM module comprised of PAS-GAF-PHY was modeled in the monomeric state (apo/holo), along with the heteroatoms using the solved structures of *A. thaliana* (PDB ID: 4OUR), *G. max* (PDB ID: 6TL4) (Nagano et al., 2020) and *P. aeruginosa* (PDB ID: 3C2W) (Yang et al., 2008) in UCSF chimera (Pettersen et al., 2004), to intact the local secondary structure and ensure similar domains orientation. The PΦB was docked into the binding pocket of holo-PSM-PhyB using MOE (Inc., 2016). Next, OPM module comprising of PAS1, PAS2 and HKR domains was added to the C-terminal domain of PSM module by using multiple solved structures of histidine kinases from different prokaryotic organisms (PDB IDs: 3A0R, 4R3A and 4GCZ) (Yamada et al., 2009; Diensthuber et al., 2013; Rivera-Cancel et al., 2014). The solved template structures helped us to retain the relative orientation of adjacent multiple domains. Fold conservation and secondary structure composition in two-domain models were achieved using the refined individual domain models as one of the templates. Sequence region that codes for the central helices/hinges to connect the PSM module with the OPM module share high identity with the templates, which helped to orient the OPM domains relative to PSM module.

It was observed that the available template structures are both in monomeric and dimeric forms, while literature reports that the Citrus PhyB probably exists in the homo-dimeric state similar to *A. thaliana*. To this context, sister chains were then assembled into the homodimer assembly in UCSF Chimera using the dimeric structure of PSM of *A. thaliana* (PDB ID: 4OUR) (Pettersen et al., 2004) and full length domain architecture model of *P. aeruginosa* (PDB ID: 3C2W) (Yang et al., 2008). PrepWizard plugin of Schrödinger 2018 (Schrödinger Release 2018_4, LLC, New York, NY, 2018) was then used to perform energy minimisation and quality assessment of initial and refined models. Model analysis and structural illustrations were generated using Pymol (DeLano, 2002).

### MD simulation investigations

To get further deep insights into dynamics of PhyB, MD simulations were performed. Since the active/holo PSM modules of *A. thaliana* (PDB ID: 4OUR) and *G. max* (PDB ID: 6TL4) (Nagano et al., 2020) are available, four different systems, including these two as references and Citrus PSM-PhyB modules in apo and holo states were explored with 100 ns MD simulations using Amber16 (Pearlman et al., 1995). The parameterisation of systems was performed with the ff14SB force field (Maier et al., 2015) and system neutralisation along with addition of missing hydrogen atoms was done by tleap module (Khalid et al., 2020). The systems were immersed in orthogonal boxes with TIP3P-model water molecules and periodic boundary conditions (Mark and Nilsson, 2001). The long-range electrostatic interactions were calculated using Particle Mesh Ewald (PME) method (Darden et al., 1999) with threshold of 10 Å. After parameterisation, 20,000 steps minimisation of the systems was performed to exclude steric clashes, and the temperature was controlled from 0 to 300 K at constant volume (NVT) for 200 ps using the Langevin thermostat during the subsequent annealing. All bonds constraints involving hydrogen atoms involved in the structures, as well as covalent bonds bearing hydrogens were restrained via SHAKE algorithm (Gonnet, 2007) and overall systems were simulated for 100 ns at 300 K and 1 atm constant pressure (NPT) using Amber16 (Pearlman et al., 1995). After every 2 fs trajectory snapshots were also obtained.

After simulations, trajectories of all the systems were evaluated by the CPPTRAJ (Amadei et al., 1993; Roe and Cheatham III, 2013). Essential dynamics and stability of the structures and systems were evaluated by calculating basic root mean square deviation (RMSD), radius of gyration (Rg) and root mean square fluctuation (RMSF). VMD (Humphrey et al., 1996) was used to visualise trajectories, whereas Xmgrace (Turner, 2005) was used to draw the plots. Furthermore, CPPTRAJ of Amber16 helped to strip the solvent and ions from the trajectories before principal component analysis (PCA) (Wood and McCARTHY, 1984), as evaluation and interpretation of the multivariate data detecting all correlated variable clusters is necessary to study essential dynamics. Diagonalisation of covariance lattices helps with figuring the central segments of movement (Van Aalten et al., 1995). The trajectories were projected over eigenvectors corresponding to the first three largest eigenvalues of the correlation matrix. Then NMWiz VMD plugin was used to investigate the most promising fluctuations of different modes (Bakan et al., 2011).

### Binding free energy calculations

The total binding free energies (ΔG_total_) of PΦB bound holo-PSM modules of *A. thaliana, G. max* and Citrus PhyB systems were calculated by using Amber16 Generalized Born and Surface Area solvation (MM/GBSA) module (Miller III et al., 2012; Genheden and Ryde, 2015) on 1000 snapshots extracted from the last 20 ns of the simulation period. Estimation of the MM/GBSA binding free energy can be concluded as:

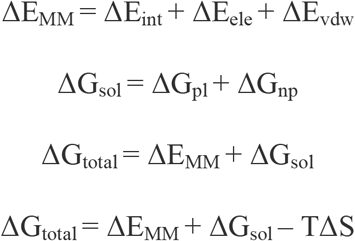

In the above equations, ΔE_MM_ is divided into internal energy (ΔE_int_), electrostatic energy (ΔE_ele_), and van der Waals energy (ΔE_vdw_), and the polar (ΔG_p_) and non-polar (ΔG_np_) energy components contributed to total solvation free energy (ΔG_sol_). ΔG_total_ is the free energy of binding evaluated after entropic calculations (-TΔS), for both MM-GBSA and MM-PBSA methods. In order to estimate the decisive role of interacting residues towards ligand’s binding, per-residue energy decomposition analysis was also performed using the MM-GBSA method, and binding energy was calculated as ΔG_residue_ using the following equation.

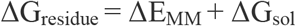

The ΔG_residue_ includes the total energy obtained from the sidechain and backbone energy decomposition. The amino acids within 8 Å of GAF domain active site were considered. The energy contributions from each residue summed up are equal to the overall binding energy of the system (Genheden and Ryde, 2015).

## ACKNOWLEDGMENTS

The authors would like to acknowledge Arooma Maryam and Rana Rehan Khalid (Department of Biosciences, COMSATS University Islamabad (CUI), Pakistan) for their kind assistance in structural modeling and simulation.

## SUPPLEMENTAL DATA

**Supplemental Figure 1.**
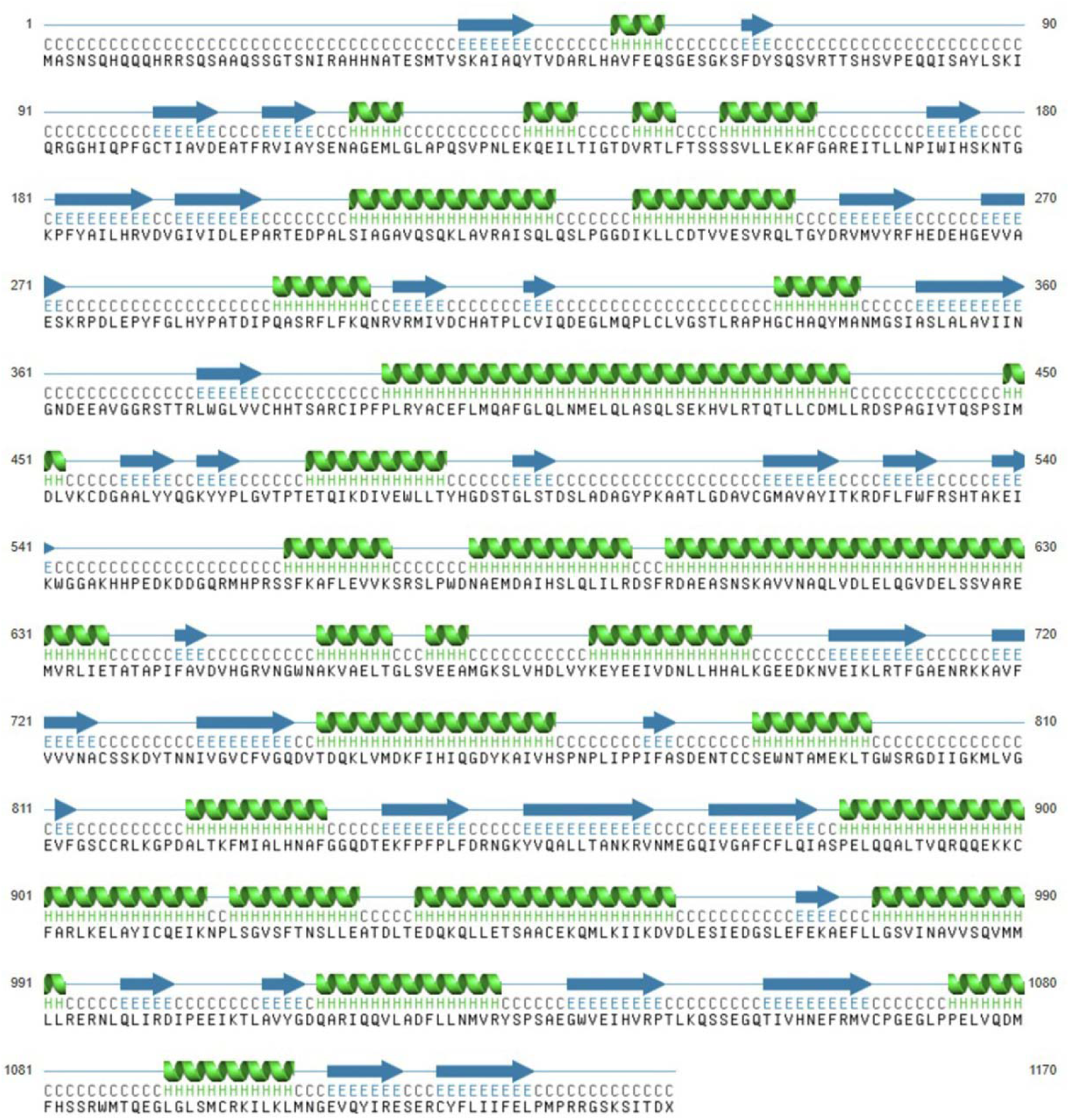
Secondary structure representation of Citrus PhyB.

**Supplemental Figure 2.**
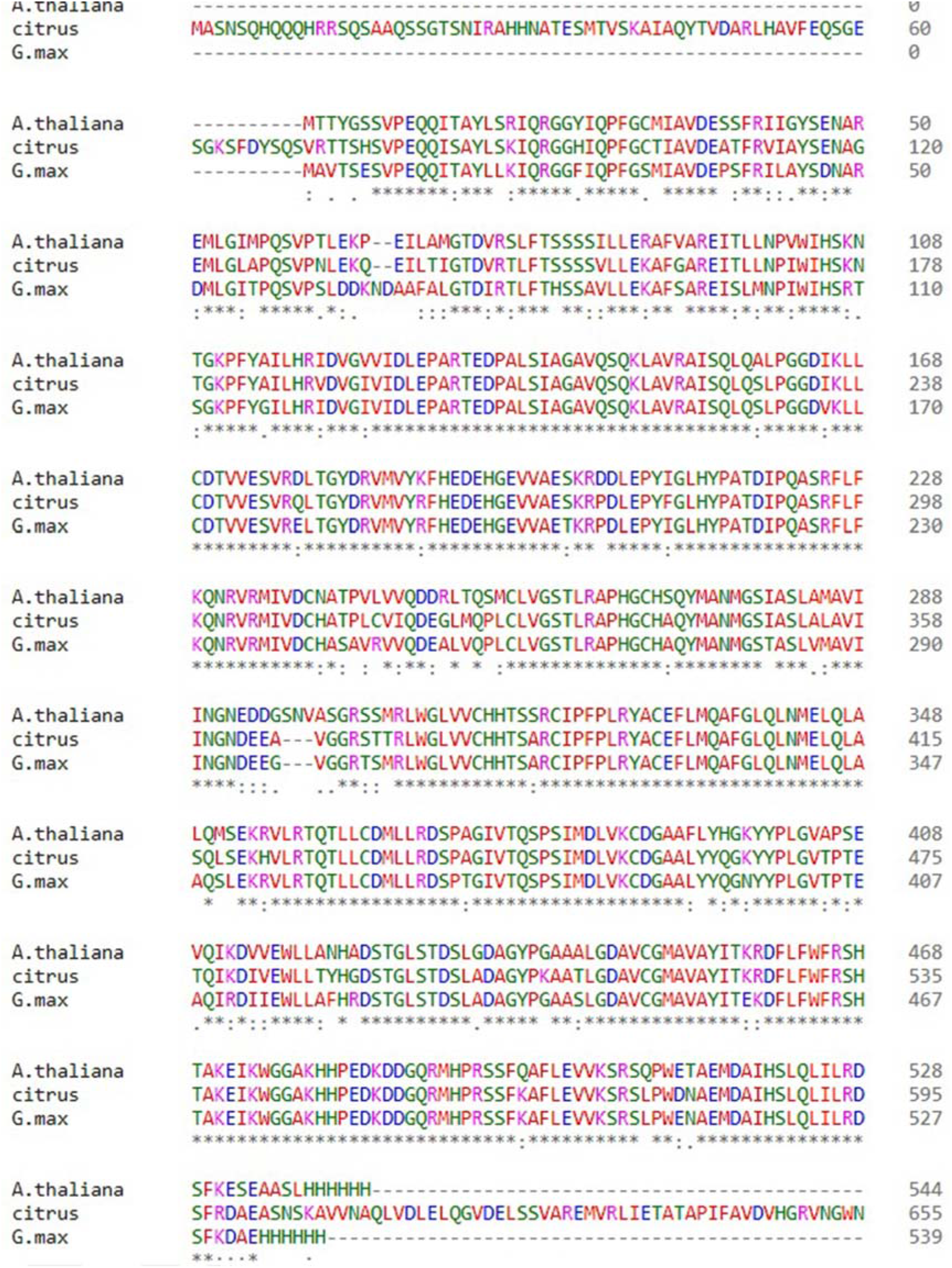
Multiple sequence alignment of Citrus, *A. thaliana* and *G. max* PSM regions.

**Supplemental Figure 3.**
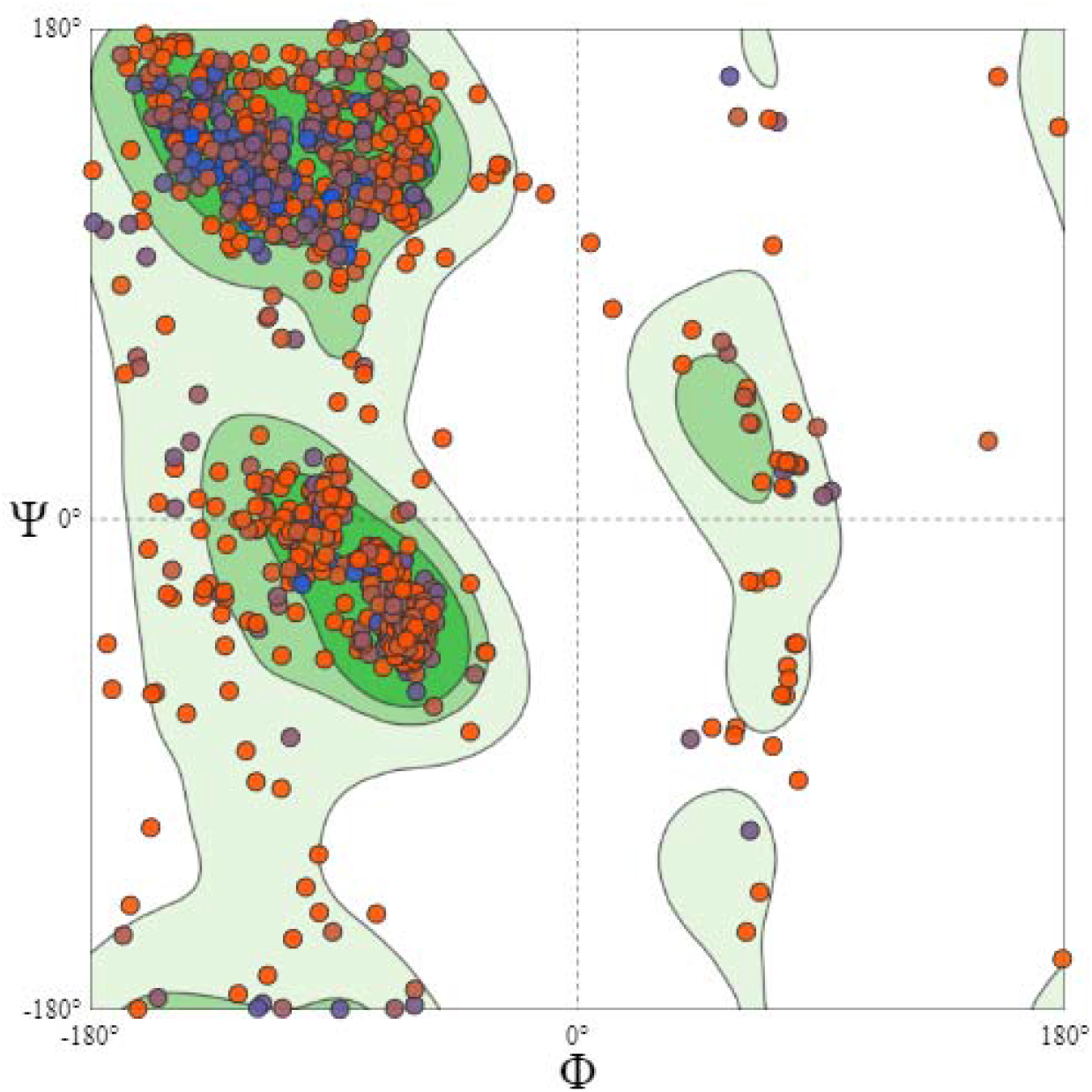
Ramachandran plot of Citrus PhyB predicted structure.

**Supplemental Figure 4.**
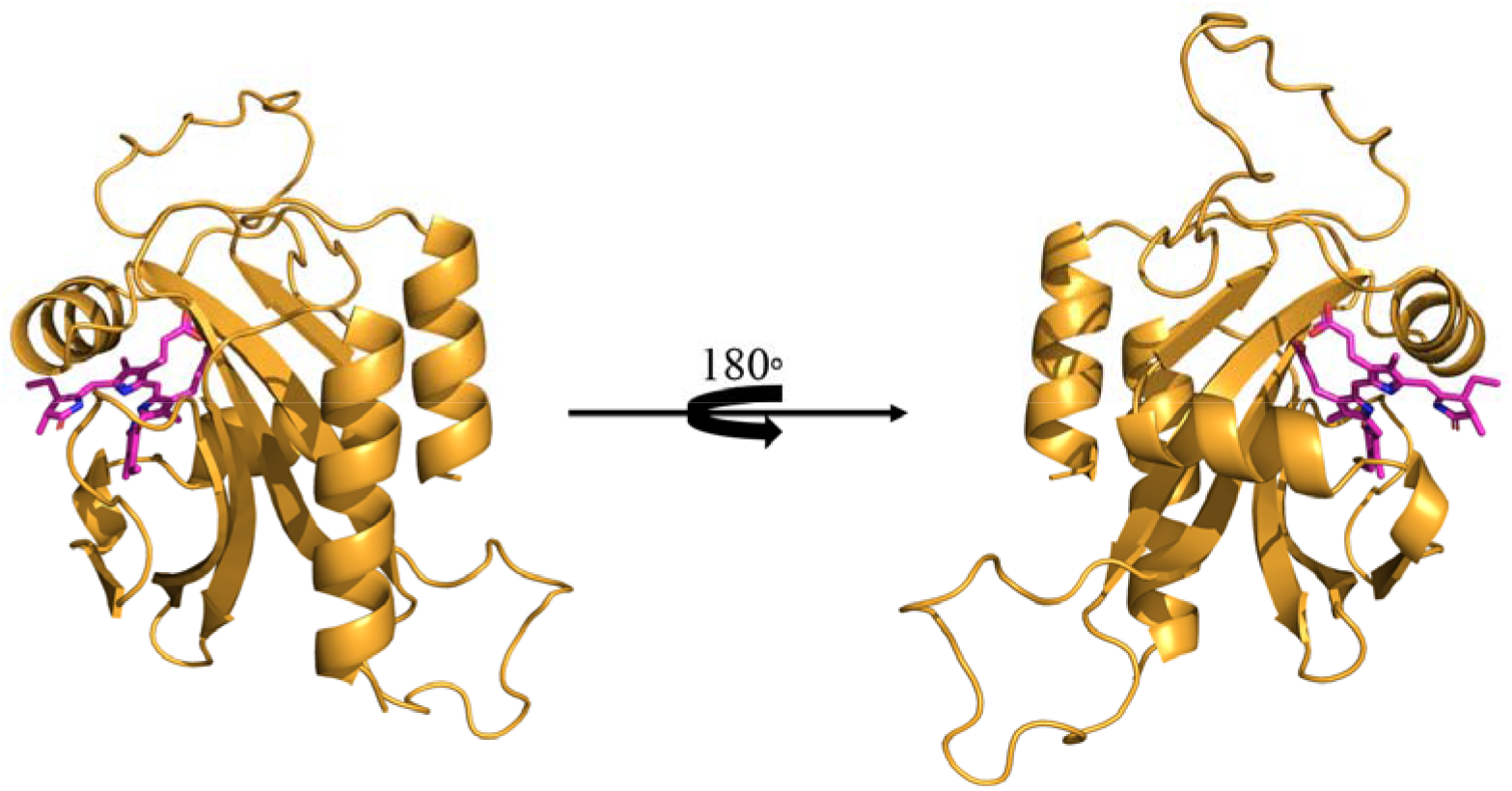
Front and back view of ligand bound 3D structures of Citrus PhyB GAF domain.

**Supplemental Table 1.**
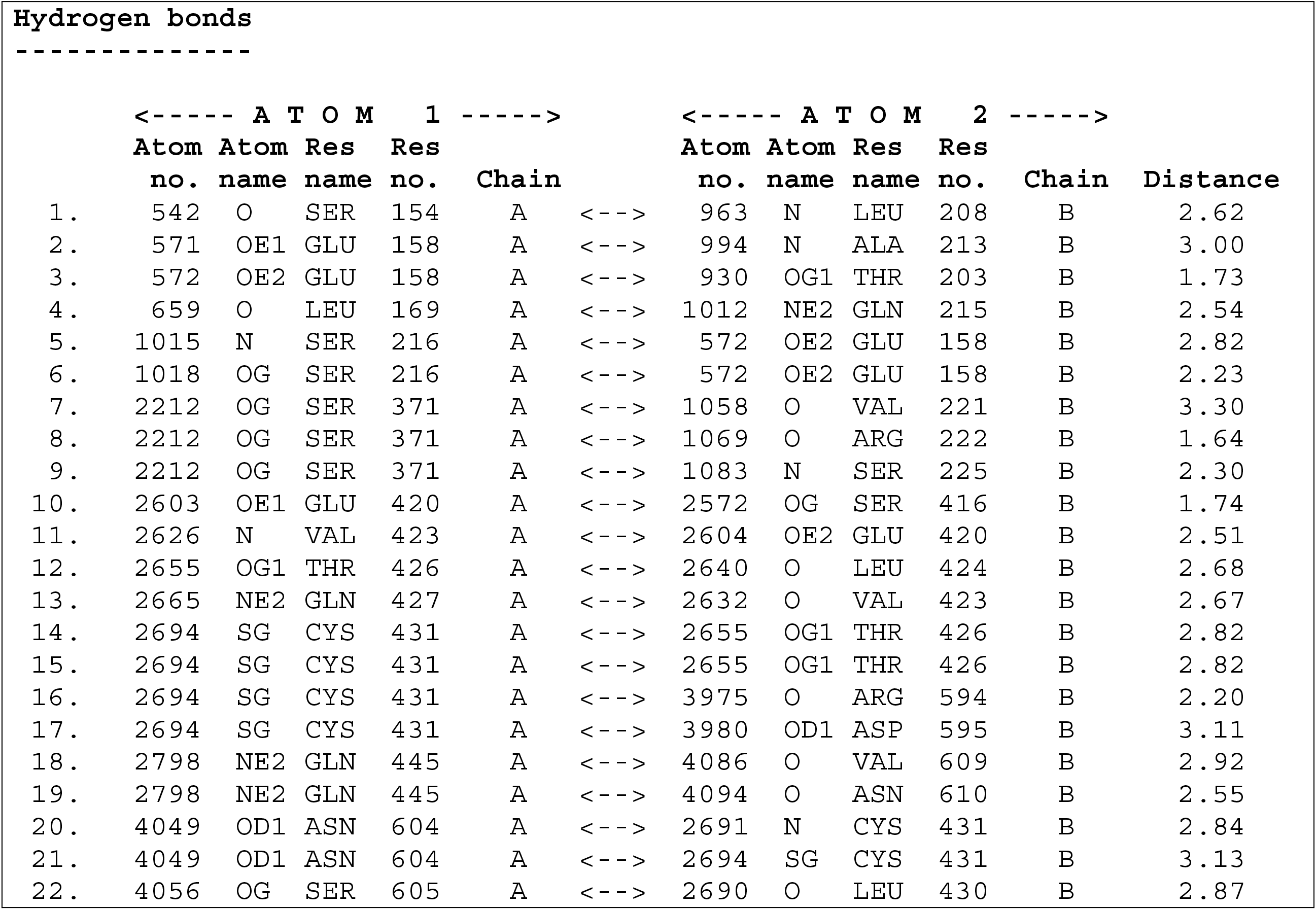
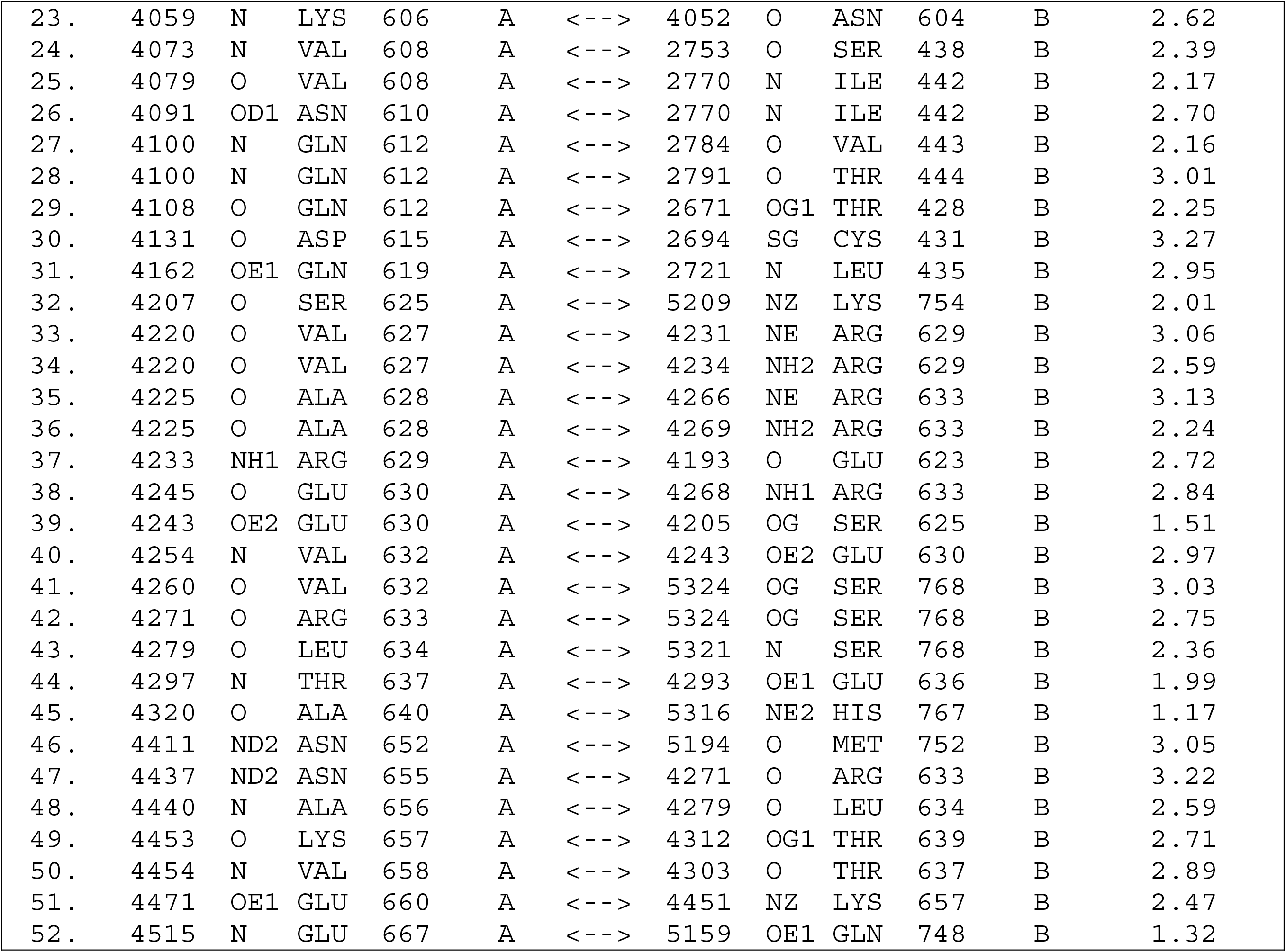
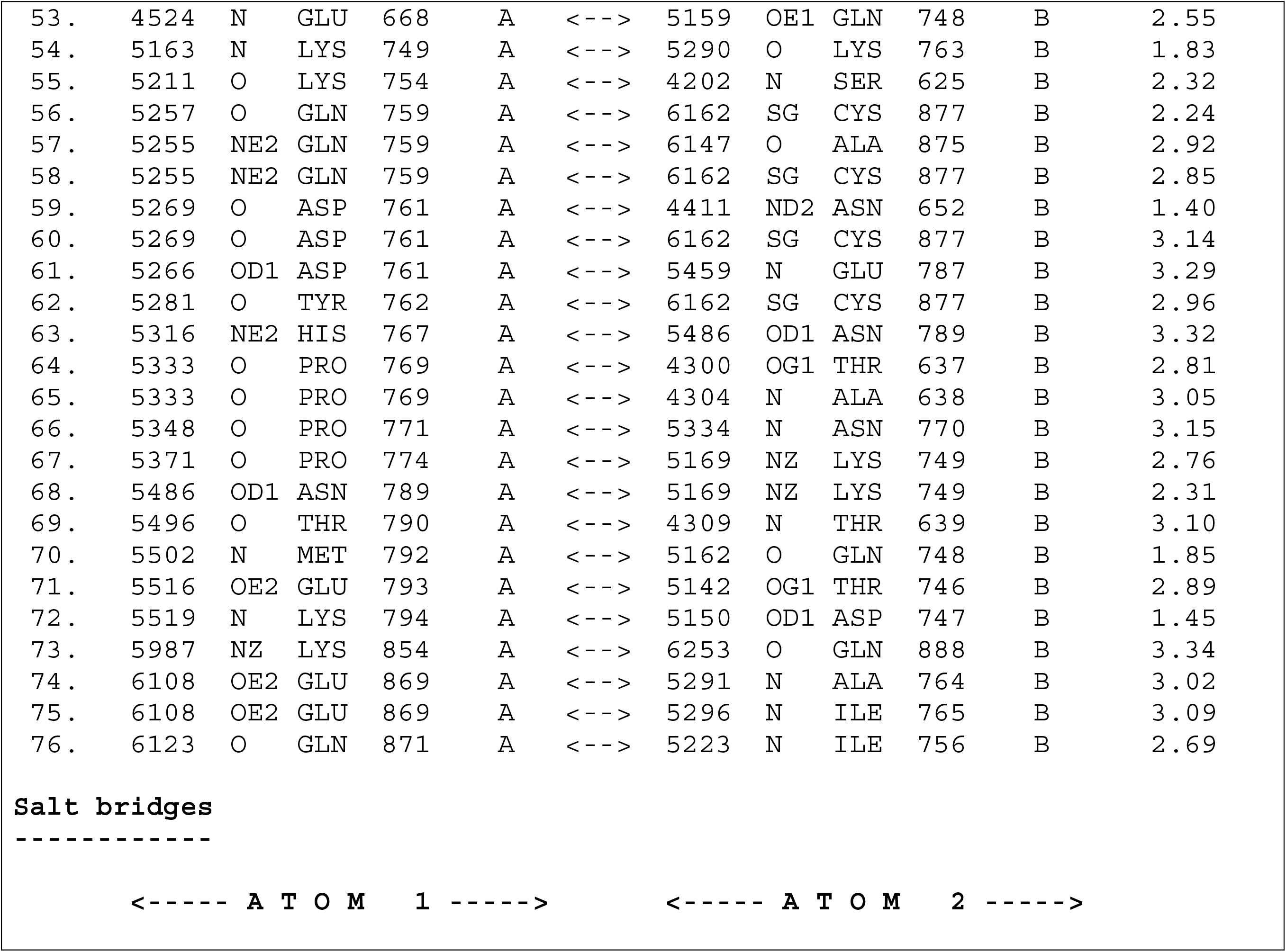
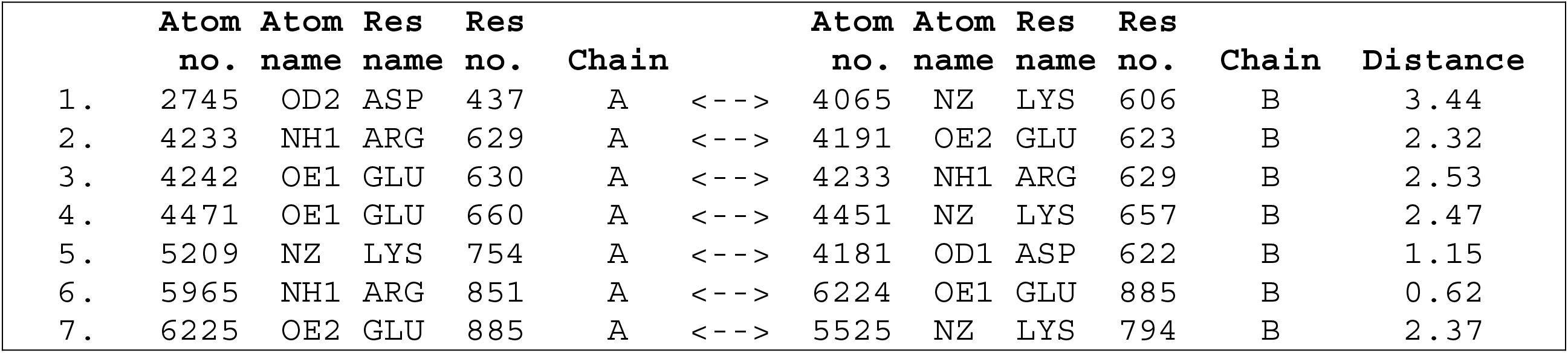
Interacting residues of both chains of Citrus PhyB homodimer.

